# Using activity data to estimate brown bear den exit and entry dates

**DOI:** 10.64898/2026.03.30.715338

**Authors:** Baptiste Brault, Jeanne Clermont, Andreas Zedrosser, Andrea Friebe, Jonas Kindberg, Fanie Pelletier

**Affiliations:** Département de biologie, Université de Sherbrooke, Sherbrooke, Québec, Canada; Department of Natural Sciences and Environmental Health, University of South-Eastern Norway, Bø, Norway; Institute of Wildlife Biology and Game Management, University of Natural Resources and Life Sciences, Vienna, Austria; Norwegian Institute for Nature Research, Trondheim, Norway; Department of Wildlife, Fish and Environmental Studies, Swedish University of Agricultural Sciences, Umeå, Sweden

**Keywords:** Phenology, Hibernation, Denning behavior, Movement, Accelerometer

## Abstract

**Background:** In hibernating mammals, the timing of den entry and exit reflects complex interactions among environment, physiology, and energetic constraints, with consequences for fitness. Consequently, shifts in denning phenology can affect population dynamics, particularly under climate change. Reliable estimation of denning timing is therefore critical, yet current methods often rely on GPS-derived movement data, limited by coarse sampling intervals, detection issues, and the inability to distinguish true inactivity from active presence at the den site. In this study, we developed and apply a method to estimate denning phenology in a brown bear population in south-central Sweden using accelerometer-derived activity data. Our approach employs adaptive, individual-specific thresholds to account for variation in baseline activity across bears, focusing on day-to-day changes to identify the start and end of inactivity periods. This method allows flexible and reproducible detection of den entry and exit dates, overcoming limitations associated with fixed thresholds and small sample sizes.

**Results:** We compared activity-based estimates with GPS-derived den occupancy and examined variation in denning behavior across demographic groups. Analyzing 388 bear-winters, the method successfully identified inactivity periods in 360 cases. The method failed to identify clear start and end dates of hibernation for 28 (7%) bear-winters, which were characterized by unusually high or low daily activity levels at the boundaries of the inactivity period. Den site occupancy ranged from September 5 to June 2, with durations of 112–260 days, whereas inactivity periods detected from activity data extended from September 6 to May 13, lasting 83–217 days. Our comparison of activity-based and GPS-based methods indicates that bears may arrive at the den site several weeks before the onset of inactivity, with timing varying among demographic groups.

**Conclusion:** We show that activity-based analysis provides a robust framework for estimating denning phenology, distinguishing actual inactivity from site presence, and improving understanding of the timing and variability of bear denning behavior. Applying an individual-level activity-based method improves accuracy in assessing ecological mechanisms underlying hibernation in bears and other hibernators, while also enhancing interpretation of environmental drivers and providing a reliable tool to monitor phenological shifts in response to climate change.

## Background

Hibernation in mammals is a seasonal physiological adaptation that allows animals to survive periods of cold temperatures and limited food availability by significantly reducing metabolic activity (1). The timing of entering and exiting a hibernation den is affected by several factors including environmental conditions, physiological state, and energetic constraints (2–4). Shifts in the duration of denning can have major impacts on body condition, reproduction, and individual survival, and ultimately on population dynamics (5–7). In the context of climate change, phenological shifts in denning behavior have been observed in several species (4,8,9). For example, as winters are getting shorter in Alaska, USA, female arctic ground squirrels (*Urocitellus parryii*) stop hibernation 10 days earlier than they used to 25 years ago (4). Accurate estimation of denning timing and long-term monitoring of changes in this important behaviour is essential for assessing the potential effects of climate change on hibernating species. Thus, reliable methods to estimate dates of key phenological events are required.

Several methods have been developed to estimate den entry and exit dates. For example, den exit can be inferred from direct visual observations of emergence events or from the presence of tracks at the hibernaculum entrance, e.g. in rodents (9,10). Other approaches include the use of antenna systems placed at the entrance of the den to record entry and exit events of individuals equipped with tags, as applied in bats (11,12), or the deployment of camera traps, as for the Eurasian badger (*Meles meles*) (13). Overall, these methods can provide relatively fine-scaled estimates of denning phenology. However, these approaches are difficult to apply to species that are sensitive to disturbance or that frequently change den sites, as den locations may vary among years and are therefore unpredictable, necessitating alternative approaches.

In hibernating bears (*Ursus arctos*, *U. thibetanus*, and *U. americanus*), although some studies have relied on observations of tracks or direct observations of bears to estimate den entry and exit dates (14,15), VHF and GPS data are by far the most commonly used sources of information (2,16–20). The advent of VHF and GPS tracking has allowed for more refined estimates, either by using the loss of VHF or GPS signals as an indicator of den entry (18–20), by identifying periods when signals become stationary (16–18), or by analyzing changes in movement speed and direction (2). Nevertheless, these approaches rely on indirect indicators of hibernation and may lack precision, as bears often reduce their movements gradually over several days or even weeks before fully retreating into their dens (16,17,21). Other studies have used body temperature thresholds recorded with implanted sensors to determine den entry and exit dates (22,23), as is sometimes done in other species, although such methods are generally applied to study hibernation physiology rather than denning phenology per se (4,13).

Activity data recorded by accelerometers mounted on collars have also been used to infer den entry and exit dates (21,24–26). An activity index as well as a threshold below which the animal is assumed to be inactive can be calculated based on acceleration values to identify prolonged periods of inactivity corresponding to hibernation (21,25–28). However, activity thresholds can be difficult to determine in wild-living animals because their validation requires direct behavioral observations, leading studies on wild bears to rely on thresholds derived from observations of captive individuals. For example, an activity threshold was defined based on observations made on two captive brown bears during their sleep and resting periods (29), and later applied to wild individuals (21,25,26). However, thresholds based on captive individuals may not necessarily reflect conditions in the wild because classification models trained on captive animals do not always transfer reliably to free-ranging individuals (30), and accelerometer readings may be influenced by external factors such as collar tightness, which can vary among individuals and across seasons (31). However, collar activity data are more informative than GPS alone for studying denning behavior because they capture fine-scale changes in movement patterns, including subtle shifts associated with pre-denning and early den exit, as well as short forays outside the den (32). As shown by Archer et al. (2025) for denning polar bears (*U. maritim*us), activity measurements closely matched camera observations of den exit, whereas GPS data provided only coarse location information and could miss brief emergence events, potentially biasing estimates of den departure timing (32).

In this study, we used data from a brown bear population in south-central Sweden to develop a method to estimate the timing of denning, defined by the start and end of the inactive period derived from accelerometer-based activity data, rather than relying solely on movement patterns inferred from GPS locations. We used day-to-day changes in activity to define adaptive, individual-based thresholds that account for inter-individual variation in baseline activity, thereby avoiding the use of fixed cut-off values. Then, we compared the estimates obtained with this activity-based method to those derived from commonly used GPS-based approaches that infer inactivity from movement data. Finally, because denning phenology in brown bears varies with sex, age and reproductive status (16,17,20), we assessed variation in denning behavior across demographic groups to evaluate whether our method yields den entry and exit chronologies consistent with those reported in previous studies using alternative methods. In addition, because denning is a phenological behavior that may be affected by temporal environmental variation, including climate change (4,9), we evaluated interannual trends to assess whether shifts over time could affect denning timing and consequently, influence comparisons with estimates reported in previous studies.

## Methods

### Study area

The study area is located in Dalarna and Gävleborg counties as well as the southern part of Jämtland County, south-central Sweden (∼61°N, 15°E). The area is covered by heavily managed boreal coniferous forest (80%), dominated by Scots pine (*Pinus sylvestris*) and Norway spruce (*Picea abies)* (33). Other main biotopes include deciduous tree stands (mainly *Betula pubescens*, *B. pendula*, *Alnus incana*, and *Sorbus aucuparia*), lakes, and bogs (33). The undergrowth vegetation is mainly composed of lichen, mosses, grasses and herbs as well as common bilberry (*Vaccinium myrtillus*), lingonberry (*V. vitis-idaea*), and black crowberry (*Empetrum hermaphroditum*) (34).

### Capture, collaring and data collection on bears

Bears were captured after den exit in April and May with capture effort directed at females accompanied by yearlings offsprings. In addition, 27 solitary individuals were captured in their dens during hibernation in February-March. See Arnemo and Evans (35) for a description of the capture and marking procedures. Overall, we equipped 148 bears (111 females, 37 males) with GPS collars fitted with accelerometers between spring 2003 and spring 2024, resulting in a total of 388 bear-winters. All capture and handling of bears have been approved by Swedish Environmental Protection Agency (NV-01278-22) and Swedish Ethical Committee on Animal Research, Uppsala (Dnr 5.8.18-21638/2023).

During captures, a premolar tooth is extracted from bears with unknown age for age determination based on cementum annuli (36). We considered individuals ≥ 1 year and <4 years as subadults, and individuals ≥4 years as adults (37). Adult females are monitored by helicopter after den exit in the spring to determine whether they are accompanied by offsprings. We classified bears into demographic groups based on their age and reproductive status. Females were retrospectively classified as pregnant during winter if cubs-of-the-year were observed with them during helicopter monitoring in spring. If no cubs-of-the-year were observed, no evidence of cubs-of-the-year presence was found at the den site, and the female was not identified as lactating at the time of capture, pregnancy status was inferred from winter activity data or from body temperature measurements recorded by physiological sensors (25,38). For the estimation of den entry data, we used seven different demographic groups based on their status observed in spring: subadult male, adult male, subadult female, solitary non-pregnant adult female, solitary pregnant female, female accompanied by cubs-of-the-year, and female accompanied by offsprings aged one year or older. For the estimation of den exit dates, we classified bears into nine groups: subadult male, adult male, subadult female, solitary adult female, females accompanied by two-year-old offsprings, females accompanied by yearlings offsprings, and three different categories of females that had given birth in the den. The latter group was divided into three categories based on whether the female was observed with her cubs-of-the-year. If cubs-of-the-year were observed, the female was classified as “female who gave birth during the winter and with cubs-of-the-year.” If no cubs-of-the-year were observed, we classified it as “female who gave birth during the winter but lost the cubs.” Finally, if it was not possible to verify the presence of cubs-of-the-year, the female was classified as “female who gave birth during the winter, but whose cubs’ fate is unknown.

Since 2003, captured bears have been fitted with GPS-GSM collars equipped with a two-axis motion sensor (Vectronic Aerospace GmbH, Berlin). Collars are programmed to collect GPS data at intervals of 30-60 minutes from April 1 to November 30 (except for 2004 and 2005, when the intervals were set to three hours), and once a day (at noon) from December 1 to March 30. The motion sensor measures raw acceleration in units of gravitational acceleration (9.8 m/s^2^) six to eight times per second in two orthogonal directions. These values are then averaged for five-minute interval which results in mean acceleration per 5-minute interval for each axis ranging from 0 to 255.

### Statistical analysis

#### Determination of start and end dates of the inactive period

We determined start and end dates using six steps:

1. We selected all bear-winters, defined as one denning period for a given individual with activity data for each day of the potential denning period, which ranges from September until May (17).
2. We calculated a 5-minute activity index ranging from 0 to 510 by summing the mean acceleration values from the two orthogonal axes (21).
3. We used the 5-minute activity indices to calculate mean daily activity values for each individual bear-winter.
4. For each bear-winter, we carried out a Hidden Markov Model (HMM) (39) with three behavioral states (active, transitional, inactive) based on mean daily activity values. HMMs are flexible models used to analyse time series data generated by underlying hidden processes or states (40). They are commonly used to identify different behavioral states and evaluate the factors influencing transitions from one state to another (41). We used a model with three behavioral activity levels to balance model accuracy and state interpretability (42). The lowest activity level corresponds to an “inactive state”, the highest activity level to an “active state” and the intermediate level to a “transitional state”. We used a gamma distribution, because it characterizes positive continuous random variables with asymmetry near zero. We added a zero mass parameter, which combines a binomial distribution with the gamma distribution to account for the inflation of zero counts (43). Since we did not expect that the active and transitional states exhibit zero activity, we constrained the zero-mass parameter to zero for these two states, which improved model likelihood. We fitted HMMs using numerical optimisation of the log-likelihood function with the momentuHMM package (43) in the R statistical environment (44). This procedure required initial starting values for each parameter that were determined based on a data visualisation. Instead of trying only one set of starting values, we investigated a grid of plausible values to finally select the best model fit corresponding to the model with the smallest negative log-likelihood. Once the HMM parameters were estimated, we obtained for each daily observation a probability associated with each of the three states. Based on these probabilities, we calculated the most plausible sequence of states using the Viterbi algorithm (45). For each observation, we then identified the start and end dates of the inactivity period based on HMM results. The start and end dates of the inactivity period correspond to the first and last day of inactivity, respectively. We determined the duration of the inactivity period as to the number of days between the start date of inactivity and the end date of inactivity. For bears captured during hibernation in their den, we excluded the end date of the inactive period from analyses because den capture commonly results in den abandonment and thus in atypical hibernation activity patterns (46).
5. For each bear-winter, we generated a plot of mean daily activity over time. We used graphical visualization to verify that the start and end of the period of inactivity had been correctly identified by the HMM, and excluded erroneous observations.
6. We extracted start and end dates of inactivity for each bear-winters. Because captures that occurred during the denning period could influence den exit behavior, we excluded the end-of-inactivity dates from the affected observations.

Based on previously published methodology in Friebe et al. (25) and Sahlén et al. (21), we used the mean daily activity as a primary approach but we also applied the same methodology (steps 3-6) using daily activity variance. Due to very similar results, we only present results based on the mean, while results using the variance are presented in Additional file 1.

#### Determination of dates of arrival and departure from den site

We identified the location of each den in every bear-winters using GPS data to determine the arrival and departure dates of an individual bear from the den site. Because bears may occasionally switch dens during winter (21,47), our first step was to identify the location of den sites used at both the start and the end of the inactivity period identified by activity-based HMMs. Since GPS position intervals can be irregular, we applied the *crawlWrap* function from the *momentuHMM* package to interpolate temporally regular daily locations (43). We then performed a sequential clustering of the GPS data with the GPSeqClus package (48). We aggregated locations into spatio-temporal clusters based on three criteria: a 50 m search radius, a 3-day temporal window, and a minimum of three GPS locations. For each cluster identified as a den, we calculated centroid coordinates using the median of x and y coordinates and added a 50m buffer to define a den site (21).

To minimize the risk of misclassifying temporary movements, we applied strict criteria to define arrival and departure dates. The arrival date at the den site was defined as the last day of a period of at least seven consecutive days spent outside the 50m den site buffer prior to the inactivity period, a threshold chosen arbitrarily to ensure sustained site use rather than transient visits. The departure date was defined as the first day of a period of at least seven consecutive days spent outside the den site buffer following the inactivity period.

Lastly, we studied the sequential occurrence of the start and end dates of the inactivity period identified using HMMs and the arrival and departure dates at the den site estimated from GPS locations using descriptive statistics. The correlation between arrival date and start of inactivity period and between departure date and end of inactivity period was assessed using a Pearson linear correlation coefficient with its 95% confidence.

#### Variation in denning behavior across demographic groups and years

We compared start and end dates of the inactivity period across demographic groups by calculating median dates and through graphical representations. To assess whether denning phenology differed among demographic groups, we used linear mixed-effects models fitted with the lmerTest (49) package in R (44). Separate models were built for start and end dates of the inactivity period (expressed as Julian day). In each model, demographic group and year (centered on the mean sampling year) were included as fixed effects. Individual identity was included as random intercepts to account for repeated measures of the same individuals across years. Models were fitted using restricted maximum likelihood (REML). Model assumptions were assessed by visually inspecting residual plots and Q–Q plots to verify homoscedasticity, normality of residuals, and the absence of strong outliers or influential points. Significance was set at α = 0.05 for all tests.

Estimated marginal means (LS-means) were computed for each demographic group using the *emmeans* package (50). LS-means provide adjusted group means that account for the random effects structure of the model and potential unbalanced sample sizes. Pairwise contrasts between demographic groups were then performed using Tukey’s post hoc tests to control for Type I error inflation associated with multiple comparisons.

## Results

### Determination of start and end dates of the inactive period

Our visual validation showed that the HMM-based method for identifying the start and the last date of inactivity was able to identify start and end dates in 93% of the cases (360 bear-winters). We therefore concluded that the method efficiently identified the period of inactivity (Figure 1). The method could not identify clear start and end dates of the inactivity period for 28 (7%) bear-winters. These bear-winters were characterized by very high mean daily activity levels at the beginning or end of the inactivity period compared with the rest of the winter, or by similarly low activity levels immediately before or after the period of inactivity (Some examples are presented in Additional file 2).

**Figure 1.**
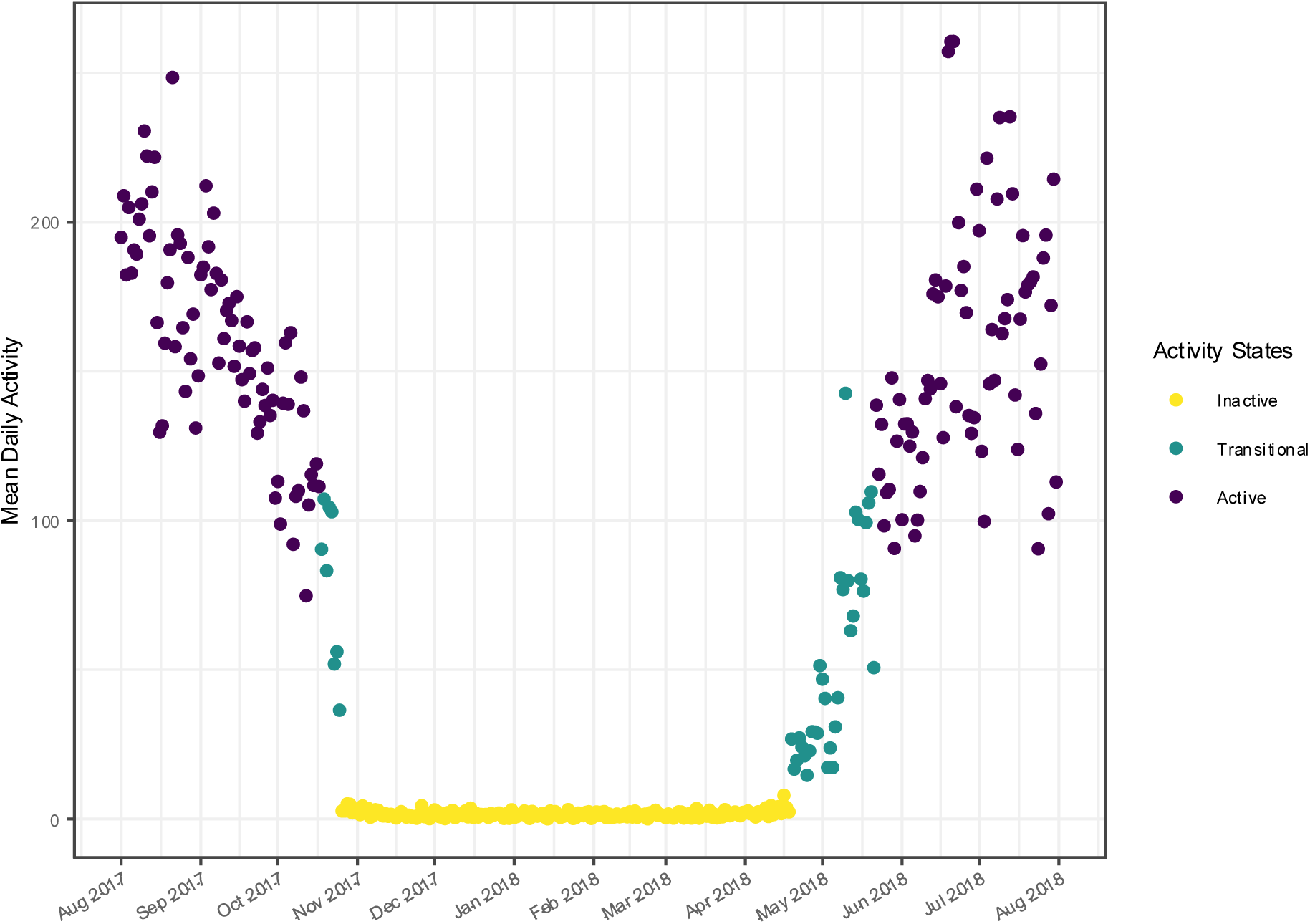
Mean daily activity and HMM states in a denning female Scandinavian brown bear. Example of mean daily activity over time for a six-year-old female Scandinavian brown bear denning with her three cubs between August 2017 and July 2018. The yellow, green, and purple colors represent the three activity states (inactive, transitional, and active, respectively) assigned by the Hidden Markov Model. The first and last days of the inactive period (yellow) were used to determine the start and end dates of inactivity.

### Discrepancy between GPS and activity-based methods

Out of 360 bear-winters, GPS data enabled the identification of the first den used in 75% of cases (269 bear-winters). For the estimation of den exit dates, we excluded 27 bear-winters because the bears were captured during hibernation, which may have influenced the timing of den exit. Among the remaining 333 bear-winters, GPS data allowed identification of the last den used in 57% of cases (190 bear-winters). In 268 bear-winters, the arrival date at the den site coincided with or preceded the start of inactivity (Figure 2A). All 190 departure dates from the den site occurred after the end of inactivity (Figure 2B).

**Figure 2.**
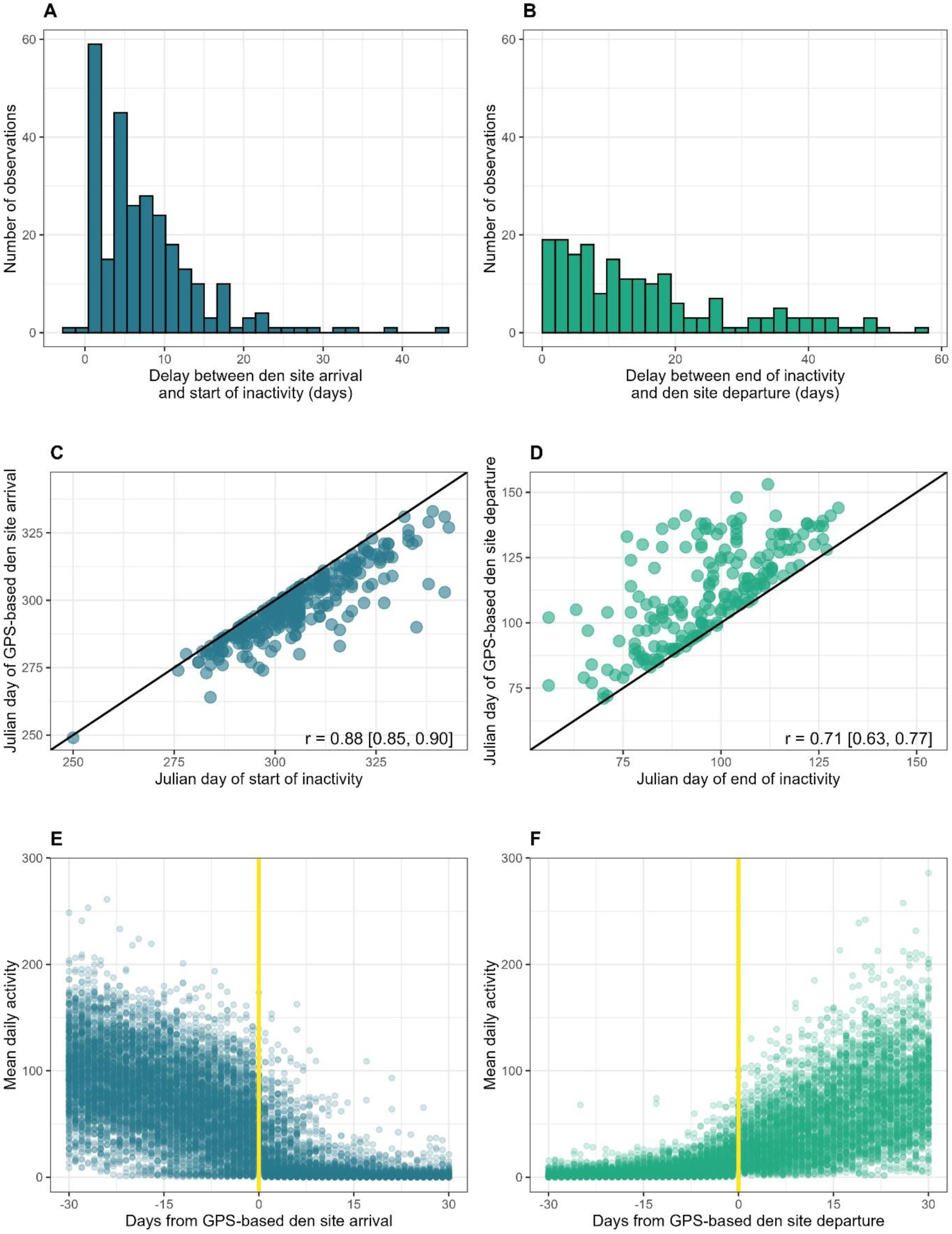
Timing of den site use and inactivity periods in Scandinavian brown bears. A) Number of bear-winters as a function of the number of days between arrival at the denning site and the start of the inactivity period, and B) between the end of the inactivity period and departure from the denning site of brown bears in south-central Sweden, 2003-2024. Correlations between C) the date of arrival at the den site and the start of the inactivity period, and D) between the date of departure from the den site and the end of the inactivity period. The black line indicates the 1:1 (identity) line (y = x). The relationship between E) mean daily activity of individuals and the dates of den site arrival and F) departure indicated by time zero. The data have been aligned to the date of arrival (E) and departure (F) indicated by a yellow vertical line. The points indicate the daily average value for each of 360 bear-winters.

The correlations between den site arrival dates and start date of inactivity (Figure 2C) and between inactivity end dates and departure dates from the den site (Figure 2D) were strong and positive. In many cases, daily mean activity remained high for several days after the arrival date (Figure 2E), whereas the reverse was observed for den site departure (Figure 2F). Overall, the delay between den-site arrival and the start of inactivity were shorter than those between the end of inactivity and actual departure from the den. The mean lag between den-site arrival and the start of inactivity was 7.7 days (median = 6 days), whereas the mean lag between the end of inactivity and departure from the den site was 15.2 days (median = 11.5 days) (Figures 2; Tables 1 and 2).

**Table 1.** Timing of den site arrival and inactivity start by bear demographic group. Arrival dates at the den site, start dates of the inactivity period, and number of days elapsed between these two events for different demographic groups of Scandinavian brown bears between 2003 and 2023.

| Demographic group before the denning period | Start of inactive period |  |  | Den site arrival |  |  | Duration between arrival date and inactive period (days) |  |  |
| --- | --- | --- | --- | --- | --- | --- | --- | --- | --- |
|  | Min | Median | Max | Min | Median | Max | Min | Median | Max |
| Pregnant female | 06 Sept. | 23 Oct. | 24 Nov. | 05 Sept. | 17 Oct. | 16 Nov. | 1 | 5.0 | 28 |
| Female with cubs-of-the-year | 15 Oct. | 31 Oct. | 22 Nov. | 13 Oct. | 27 Oct. | 14 Nov. | 0 | 6.0 | 19 |
| Female with yearlings | 10 Oct. | 07 Nov. | 18 Nov. | 09 Oct. | 01 Nov. | 09 Nov. | 1 | 6.5 | 14 |
| Solitary adult female | 11 Oct. | 31 Oct. | 29 Nov. | 30 Sept. | 23 Oct. | 16 Nov. | 2 | 6.0 | 26 |
| Adult male | 15 Oct. | 07 Nov. | 04 Dec. | 07 Oct. | 02 Nov. | 28 Nov. | 1 | 6.0 | 18 |
| Solitary subadult female | 16 Oct. | 11 Nov. | 08 Dec. | 12 Oct. | 30 Oct. | 26 Nov. | 1 | 8.5 | 45 |
| Solitary subadult male | 17 Oct. | 08 Nov. | 03 Dec. | 09 Oct. | 23 Oct. | 21 Nov. | 5 | 11.0 | 33 |

Abbreviations: Min. = Minimum; Max. = Maximum

**Table 2.** Timing of inactivity end, den site departure and duration by bear demographic group. End dates of the inactivity period and start dates of departure from the den site, number of days elapsed between these two events, duration of the inactivity period, and duration of den site occupancy for different demographic groups of Scandinavian brown bears between 2003 and 2024.

| Demographic group after the denning period | End of inactive period |  |  | Den site departure |  |  | Duration between end of inactive period and departure (days) |  |  | Inactive period (days) |  |  | Den site occupancy (days) |  |  |
| --- | --- | --- | --- | --- | --- | --- | --- | --- | --- | --- | --- | --- | --- | --- | --- |
|  | Min. | Med. | Max. | Min. | Med. | Max. | Min. | Med. | Max. | Min. | Med. | Max. | Min. | Med. | Max. |
| FGB COY | 17 Mar. | 15 Apr. | 13 May | 15 Apr. | 12 May | 02 Jun. | 2 | 20.0 | 57 | 121 | 175 | 217 | 175 | 208.5 | 233 |
| FGB CL | 25 Feb. | 28 Mar. | 15 Apr. | 17 Mar. | 03 Apr. | 21 May | 1 | 12.0 | 50 | 127 | 151 | 209 | 155 | 156 | 260 |
| FGB CU | 14 Mar. | 08 Apr. | 27 Apr. | 21 Mar. | 24 Apr. | 09 May | 1 | 6.5 | 34 | 126 | 166.5 | 200 | 155 | 184 | 211 |
| F Y | 25 Feb. | 10 Apr. | 07 May | 28 Mar. | 18 Apr. | 10 May | 1 | 6.0 | 46 | 100 | 159 | 196 | 142 | 175 | 203 |
| F TYO | 15 Mar. | 04 Apr. | 23 Apr. | 03 Apr. | 14 Apr. | 04 May | 1 | 8.0 | 47 | 123 | 151 | 196 | 152 | 174 | 201 |
| AF | 07 Mar. | 06 Apr. | 23 Apr. | 23 Mar. | 13 Apr. | 02 May | 1 | 13.5 | 31 | 117 | 153 | 188 | 146 | 177 | 206 |
| AM | 04 Mar. | 24 Mar. | 09 Apr. | 13 Mar. | 31 Mar. | 13 Apr. | 1 | 4.0 | 18 | 113 | 139.5 | 171 | 132 | 153 | 174 |
| SF | 26 Feb. | 30 Mar. | 18 Apr. | 12 Mar. | 11 Apr. | 24 Apr. | 1 | 6.0 | 31 | 83 | 142 | 172 | 112 | 162.5 | 185 |
| SM | 16<br>Mar. | 02<br>Apr. | 10<br>Apr. | 20<br>Mar. | 02<br>Apr. | 15<br>Apr. | 4 | 4.5 | 5 | 109 | 143.5 | 168 | 122 | 134 | 146 |

Min. = Minimum; Med. = Medium; Max. = Maximum

### Variation in denning behavior across demographic groups and years

The start dates of the inactivity period ranged from September 6 to December 8, and the end dates ranged from February 25 to May 13, resulting in total denning duration spanning from 83 to 217 days (Tables 1, 2). The linear mixed-effects model revealed significant differences in the timing of denning activity among demographic groups. Model estimates revealed that pregnant females started their inactivity period significantly earlier, 9.9 to 17.2 days, than other demographic groups, with the exception of solitary adult females, for which no statistically significant difference was detected after Tukey adjustment (−6.9 days [95% CI: −13.81; 0.02]; p-value = 0.051). This near-significant result may reflect the conservative nature of the Tukey adjustment for multiple comparisons and the limited sample size in the solitary female category, rather than a true absence of difference. Subadult females also began their inactivity period 7.2 and 10.3 days later than females with cubs-of-the-year and solitary adult females, respectively. Model estimates showed that females giving birth during hibernation ended their inactivity 11.2 to 21.5 days later than all other demographic groups. The only exceptions were females that gave birth during winter with unknown cubs fate and females with yearlings offsprings, for whom the differences were not statistically significant. Females with yearlings offsprings ended their inactivity 9.5, 13.8, and 16.6 days later than subadult females, females that had lost their cubs during winter, and adult males, respectively. For additional details on model estimates and significance levels for comparisons between demographic groups, see Additional file 3.

Compared to other demographic groups, females with cubs-of-the-year stayed the longest at the den site after activity resumed, for a median duration of 20 days (first quartile: 13 days; third quartile: 35 days, Table 2). For females accompanied by cubs-of-the-year at den exit, the mean and median durations of the inactivity period were 174.8 and 175 days, respectively, with a first quartile 163.8 days and a third quartile 189 days (Figure 3, Table 2). In comparison, individuals that did not give birth during the denning period exhibited shorter inactivity durations, with a mean and median of 147.6 and 149 days, respectively, ranging from 135 days (first quartile) to 160 days (third quartile). Regarding total occupation time of the den site, period between arrival and departure from the site, females that gave birth and emerged with newborns stayed longer with a mean and median of 206.9 and 208.5 days (Q1-Q3: 193.8-218.3 days), respectively. In contrast, individuals that did not give birth had a mean and median of 165.9 and 167 days (Q1-Q3: 153-178 days), respectively. We found a significant effect of year on the end of the inactivity period, with bears emerging progressively earlier over the last two decades (β = −0.35 ± 0.15 days per year, p-value = 0.019), indicating a shift in denning phenology across years. In contrast, no significant effect of year was detected for the start of the inactivity period (Figure 4).

**Figure 3.**
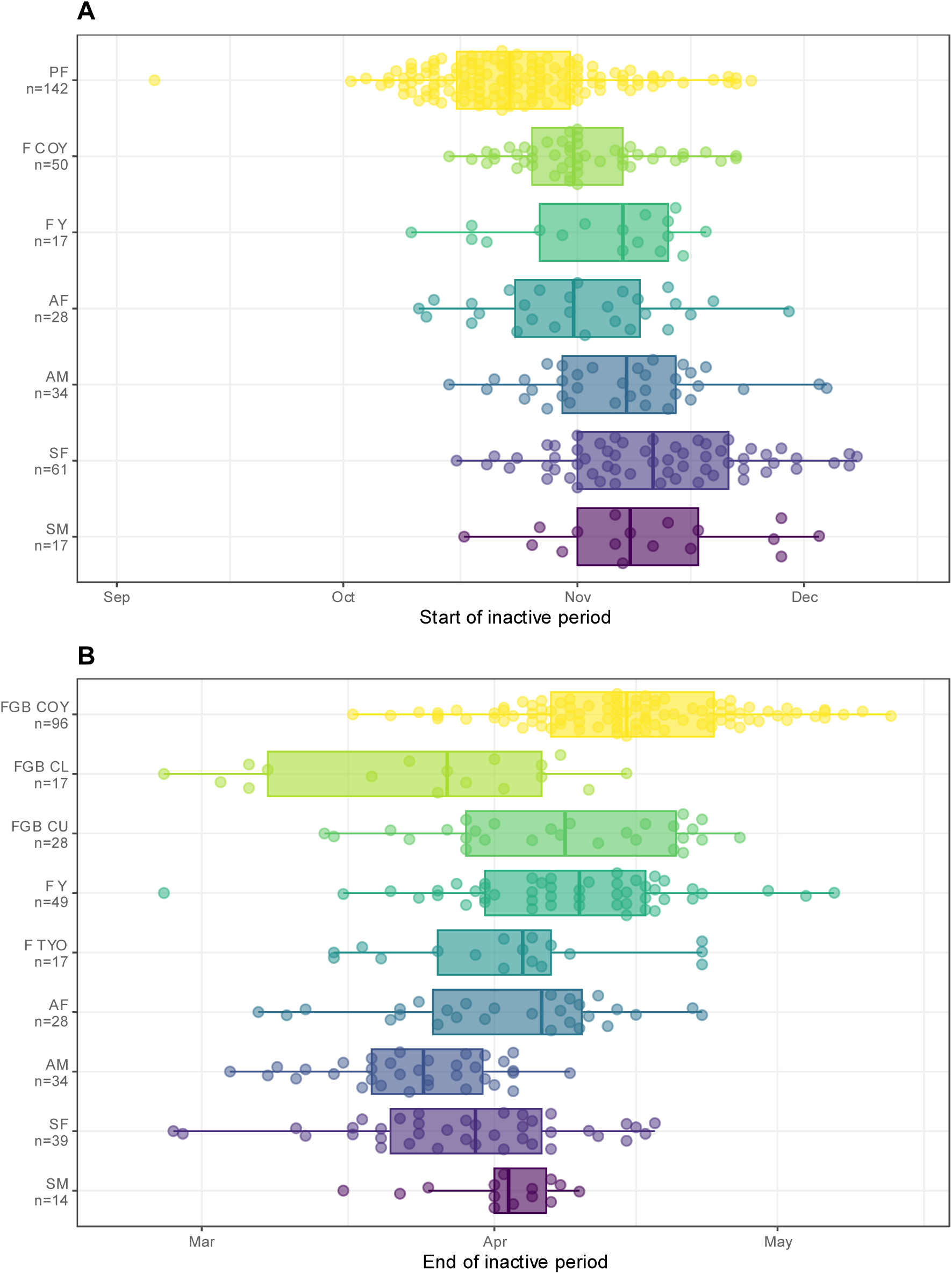
Start and end dates of brown bear inactivity periods by demographic group. Boxplots of start (A) and end (B) dates of the inactivity period by demographic group of brown bears in south-central Sweden between 2003 and 2024. PF = Pregnant female; F COY a= Female with cubs-of-the-year; F Y = Female with yearlings; F TYO = Female with two years old offsprings; AF = Solitary adult female; AM = Adult male; SF = Solitary subadult female; SM = Solitary subadult male; FGB COY = Female who gave birth during the winter and with cubs-of-the-year; FGB CL = Female who gave birth during the winter but lost the cubs; FGB CU = Female who gave birth during the winter but where the fate of the cubs is unknow.

**Figure 4.**
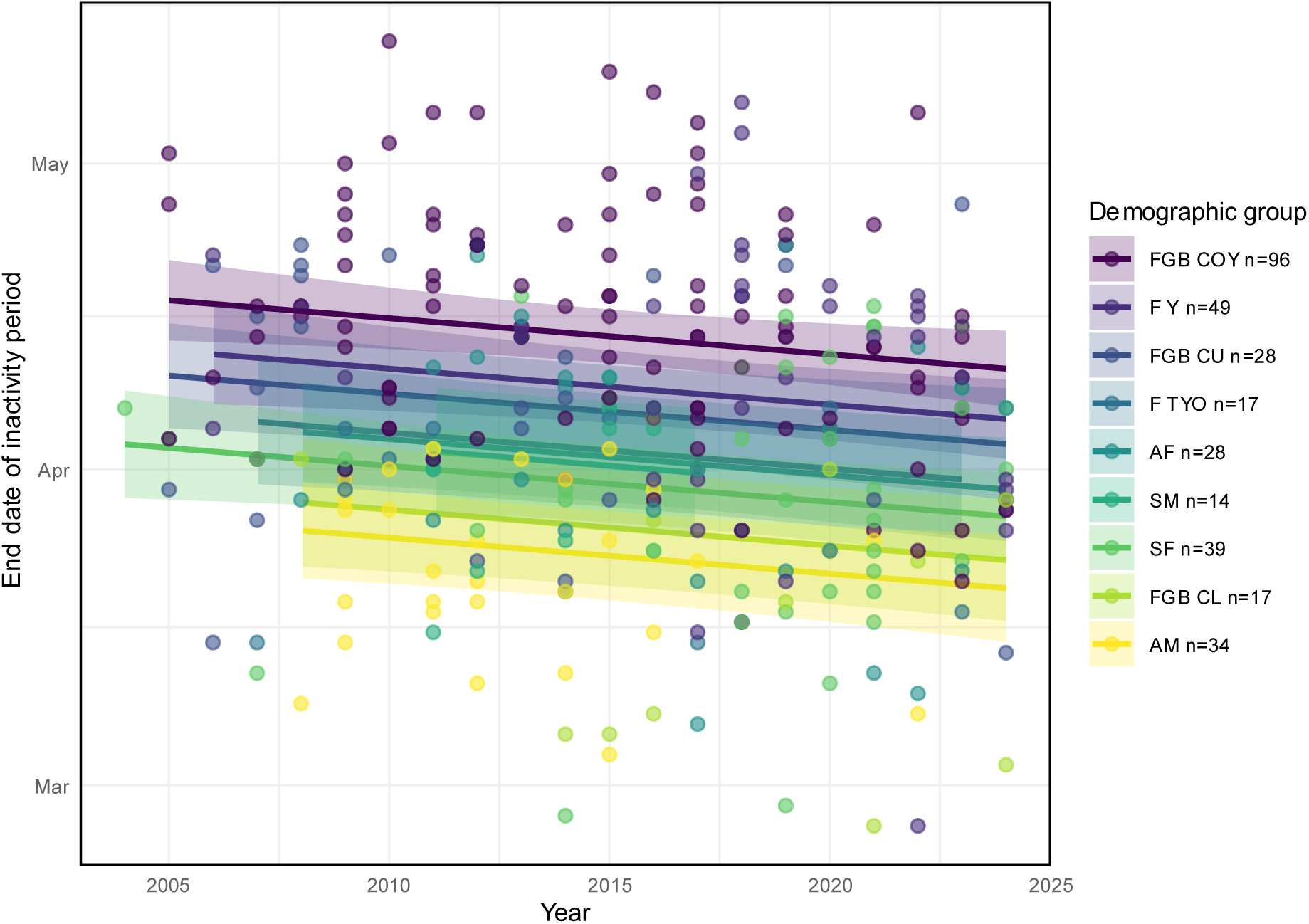
Temporal trends in inactivity end dates across brown bear demographic groups. Effect of year on the end dates of the inactivity period in brown bears by demographic group in south-central Sweden, 2004–2024. Points represent individual observations of inactivity end dates for each bear-winter. Lines show the estimated slope of the relationship between year and inactivity end date for each demographic group, with shaded areas indicating 95% confidence intervals. Demographic groups are indicated by different colors. F Y = Female with yearlings; F TYO = Female with two years old offsprings; AF = Solitary adult female; AM = Adult male; SF = Solitary subadult female; SM = Solitary subadult male; FGB COY = Female who gave birth during the winter and with cubs-of-the-year; FGB CL = Female who gave birth during the winter but lost the cubs; FGB CU = Female who gave birth during the winter but where the fate of the cubs is unknow.

## Discussion

In this study, we develop and test a flexible and reproducible framework for identifying the onset and termination of inactivity, as well as estimating den entry and exit dates. The HMM-based method successfully identified periods of inactivity for 360 out of 388 bear-winters. Among the 28 cases where classification failed, one involved a pregnant female, for whom elevated activity levels during gestation were misclassified as a transition period rather than as part of the inactivity period. Notably, this issue did not occur in the remaining 144 bear-winters of pregnant females included in the analysis. We compared these dates obtained from accelerometer-derived activity data with the dates of arrival at and departure from the den site derived from GPS data and observed a significant correlation between the two sets of measurements. This discrepancy indicates that the two approaches do not capture the same behavioral phenomenon, and that den site occupancy includes active phases that may occur both before and after the actual denning period.

### Discrepancy between GPS and activity-based methods

The estimation of den entry and exit dates solely based on GPS data has several limitations. A stationary GPS position indicates only that an individual remains at the same location, however, does not necessarily represent inactivity (2,20). Moreover, the accuracy of GPS signals generally does not allow for the detection of complete immobility (2,20,25). Finally, GPS units may fail to record locations for extended periods, thereby reducing data quality and tracking precision, a limitation that frequently occurs when bears are inside their dens (20,32). These limitations are reflected in our dataset: we were able to confidently identify the first and last den site locations for only 75% and 57% of cases, respectively, resulting in a substantial reduction in the number of den-site arrival and departure dates that could be identified. For one bear-winter, the arrival date occurred two days after the identified start of inactivity. This discrepancy likely results from the bear using its first den for fewer than three days before switching, which prevented proper identification and localization using our method.

In contrast, activity data derived from accelerometers are not subject to many of the limitations affecting geolocation systems and are typically recorded at a finer temporal resolution, providing a large number of observations per day. Furthermore, our methodology accounts for individual variability in activity patterns by operating at the individual level rather than applying a fixed activity threshold across all bears, thereby reducing bias and improving detection reliability. In addition, bear metabolism slows substantially during hibernation (2), and the space available inside dens is closely related to individual body size (51). Movements within the den are therefore constrained, likely resulting in the very low activity levels detected by accelerometers. For these reasons, we consider the start and end dates of the inactivity period to be relevant indicators of den entry and exit, respectively, whereas GPS data are more suitable for estimating general presence at a den site.

Direct comparison of the two approaches reveals a delay between the date of arrival at the den site and the start of the inactivity period, as well as between the end of the inactivity period and the date of departure from the site. In many cases, daily mean activity remained high for several days after the arrival date, whereas the opposite pattern was observed for den site departure, i.e., activity gradually increased in the weeks preceding departure, consistent with the findings of Evans et al. (2). These discrepancies were particularly pronounced at den exit, sometimes lasting several days or even weeks, which may also explain why studies do not consistently detect clear relationships between climatic factors and den entry and exit dates. For instance, Evans et al. (2), who defined den exit dates based on movement patterns detected by GPS data, found no significant climatic effect on den exit. In contrast, Pigeon et al. (20), who identified exit dates based on GPS signal recovery after total loss following entry into the den, found an effect of temperature on den exit as well as an effect of winter precipitation on den entry.

### Denning phenology in our study area

Our results indicate that bears may arrive at the den site up to several weeks before the onset of inactivity with variation among demographic group. This delay was longer for subadults than for solitary adults. This pattern is consistent with the findings of Sahlén et al. (21), who reported a decrease in the time spent at the den site prior to entry with increasing age, possibly reflecting greater experience and more efficient den construction and denning behavior in older individuals. From a management perspective, this pre-denning period may represent a critical window of increased human–bear interactions. In Sweden, many incidents involving humans, particularly hunters, occur during this period, often in close proximity to den sites (21,52). Changes in bear behavior at that time, including reduced activity and a lower propensity to flee, may contribute to this elevated risk, together with the high presence of hunters and hunting dogs (52).

Over the period 2003–2024, den entry dates ranged from October 2 to December 8, except for a pregnant female that entered her den as early as September 6. This range is slightly wider than that reported for the same study area between 1986 and 2001 based on VHF telemetry, where den entry and exit were defined as the midpoint between the last location outside the den and the first location inside the den, and den entry occurred between September 28 and November 20 (16,17).

Consistent with previous work from the same population (16), pregnant females exhibited earlier median den entry dates than other demographic groups. The only exception was solitary adult females. Beyond, the chronology among other groups appears more nuanced. For the other demographic groups, solitary adult females and females with cubs-of-the-year largely overlapped in their den entry dates, followed by subadults, adult and subadult males, and females with yearlings offsprings, which could not be clearly distinguished from each other.. Adult and subadult males, as well as females with yearlings offsprings, could not be clearly distinguished from the other demographic groups, with the exception of pregnant females. These limited differences among groups likely reflect both the relatively small separation in mean den entry dates and reduced statistical power for some categories, particularly females with yearlings offsprings and subadult males, for which sample sizes were low (17 bear-winters each).

Over the last 20 years, we observed an advancement in the end of the inactivity period, with brown bears emerging progressively earlier by approximately 7 days. This shift in denning phenology may reflect plastic responses to interannual environmental variation, such as warmer spring temperatures or changes in food availability but that remains to be assessed. Interestingly, no corresponding trend was detected for the start of the inactivity period, suggesting that there has been no long-term change in environmental drivers over these 20 years, although weather effects such as ambient temperature and snow depth, which have been shown to influence den entry in our study area (2), may still play a role. Overall, these results support the hypothesis that den exit timing is affected by environmental conditions, although the precise drivers remain to be determined. This finding is consistent with the study by Pigeon et al. (20) on Grizzly (*U. arctos horribilis*) that showed which showed that warmer spring temperatures and reduced winter snow cover triggered earlier den emergence. In addition to potential effects of climate change and local weather, other environmental changes could contribute to this advancement; for example, human activities in late winter and spring have occasionally been observed to trigger den abandonment for some bears in our study area (21,47).

## Conclusion

In conclusion, our study highlights the value of activity-based analyses for accurately identifying den entry and exit in brown bears. By distinguishing true inactivity from simple den-site presence, this approach overcomes key limitations of GPS-only methods and reveals substantial lags between arrival and den entry, as well as between den exit and final departure. These results emphasize the complexity of denning transitions and are essential for correctly interpreting links between denning phenology and environmental drivers, particularly under climate change. Overall, individual-level activity data provide a robust framework for improving phenological estimates and advancing our understanding of hibernation ecology in bears and other hibernating species.

## List of abbreviations

HMM: Hidden Marcov Model
PF: Pregnant female
F COY: Female with cubs-of-the-year
F Y: Female with yearlings
F TYO: Female with two years old offsprings
AF: Solitary adult female
AM: Adult male
SF: Solitary subadult female
SM: Solitary subadult male
FGB COY: Female who gave birth during the winter and with cubs-of-the-year
FGB CL: Female who gave birth during the winter but lost her cubs
FGB CU: Female who gave birth during the winter but whose the fate of the cubs is unknow

## Declarations

### Ethics approval and consent to participate

All capture and handling of bears have been approved by Swedish Environmental Protection Agency (NV-01278-22) and Swedish Ethical Committee on Animal Research, Uppsala (Dnr 5.8.18-21638/2023).

### Consent for publication

Not applicable.

### Availability of data and materials

The datasets and scripts supporting the conclusions of this article are available on OSF (https://doi.org/10.17605/OSF.IO/4CV8J). Raw acceleration and GPS datasets are available from the corresponding author on reasonable request.

### Authors’ contributions

BB, JC, AZ, and FP conceptualized the study and created the proposed methodology. AZ, AF, and JK participated in fieldwork and data acquisition. BB performed the analyses, interpreted the results, and wrote the manuscript. JC, AZ, FP, AZ, and JK substantively reviewed and edited the manuscript. All authors read and approved the final manuscript.

## Acknowledgement

We thank the field crew who took part in the bear captures, especially David Ahlqvist, as well as Biosphere Expeditions for their work in mapping the dens. Language editing was assisted by the free online version of ChatGPT (OpenAI, GPT-5.2). All content was subsequently reviewed and finalized by the authors.

## Additional files

**Additional file 1 - Comparison of denning dates estimated from daily mean vs. variance activity using HMM**

We applied the same methodology using Hidden Markov Model (HMM) but based on daily variance activity instead of daily mean activity (See step 3 of the methodology). We studied the correlation between den entry and exit dates obtained by HMM using daily mean activity and daily variance activity with Pearson linear correlation coefficient with its 95% confidence. The strict concordance between mean and variance was 62% for den entry and 66% for den exit, with 77% of entry dates and 79% of exit dates falling within a ±2-day agreement window. (Figure A1.1).

**Figure A1.1.**
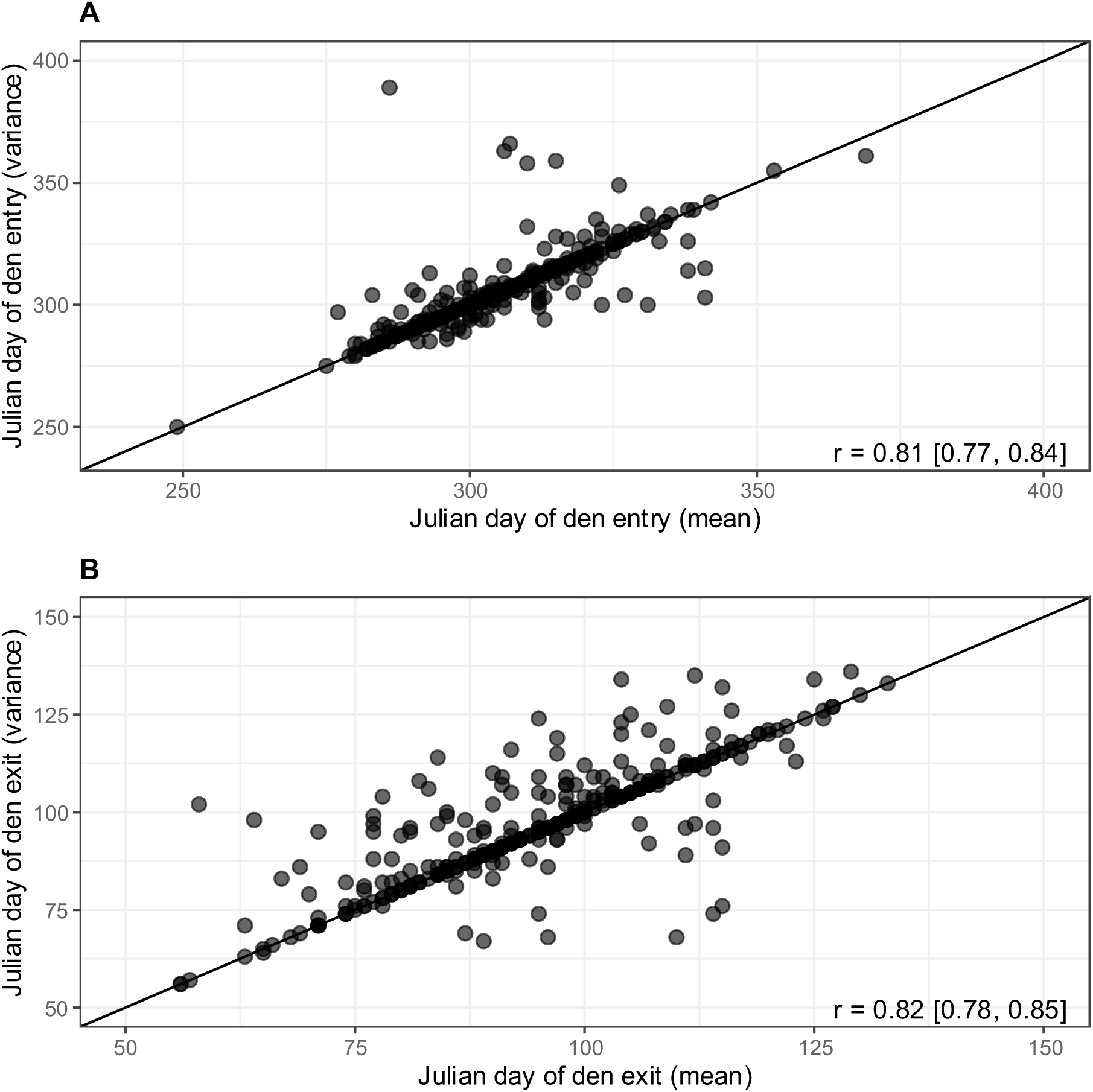
Correlation of den entry (A) and exit (B) dates obtained using Hidden Markov Model based on daily mean activity and daily variance activity.

**Additional file 2 - Visual validation and exclusion of incorrectly classified inactivity periods using HMM**

Visual validation of the HMM classifications based on daily mean activity values led us to exclude 28 observations for which the start date of the inactivity period, the end date, or both did not appear to be correctly identified. These cases were generally characterized by one of the following patterns:

- daily mean activity values at the beginning and/or end of the inactivity period classified as inactive, despite being substantially higher than activity levels observed during the rest of the winter (Figure A2.1 A and D);
- daily mean activity values immediately before and/or after the inactivity period classified as transitional, despite being similar to, or only marginally higher than, activity levels during the rest of the winter (Figure A2.1 B);
- elevated daily mean activity during gestation incorrectly classified as belonging to the transitional state (one case among the 28 excluded observations, Figure A2.1 C).

**Figure A1.1.**
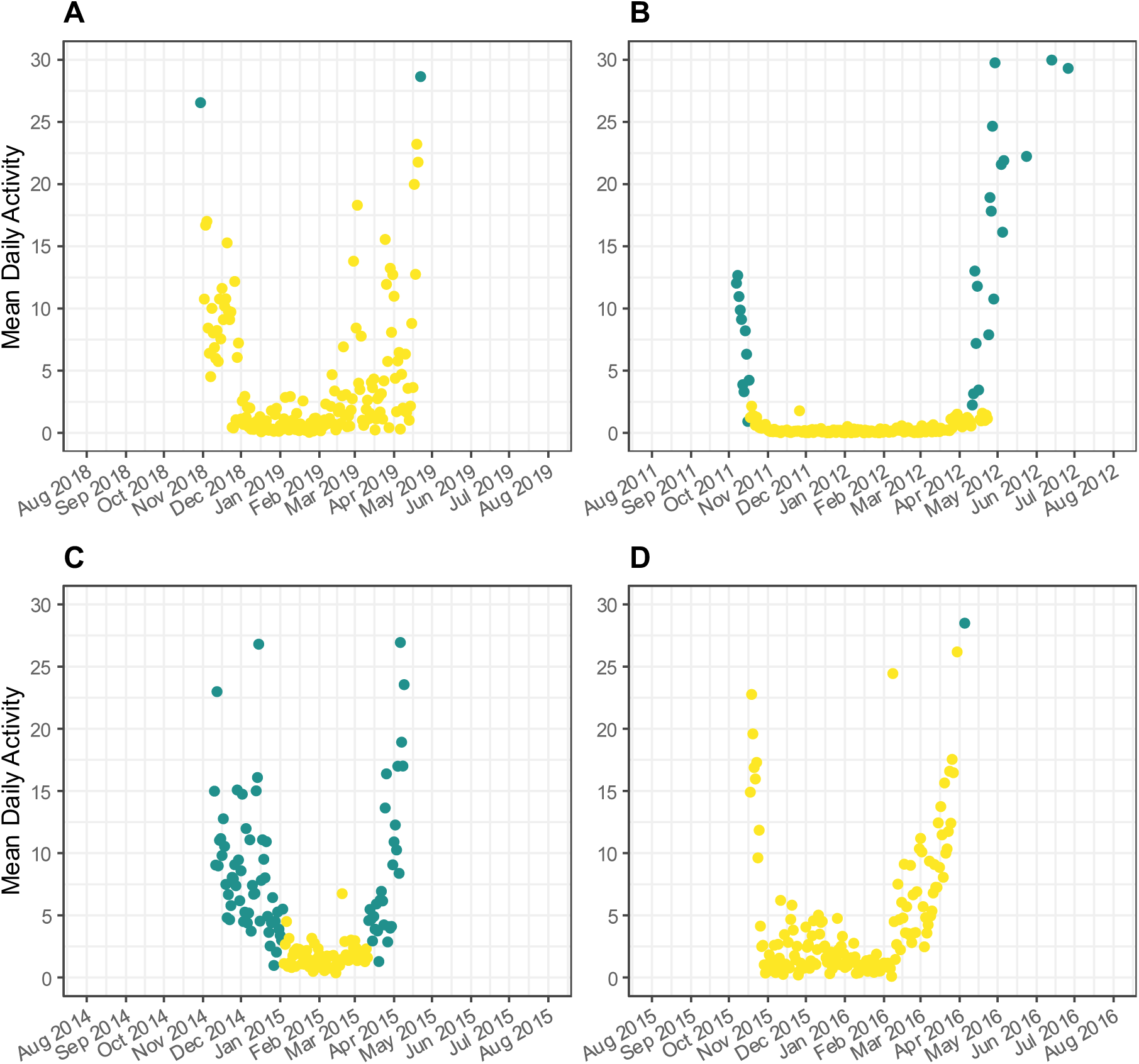
Examples of mean daily activity over time for different individuals between August and July that were excluded following visual validation of the HMM outputs. The y-axis was truncated at 30 to improve visual interpretation of transitions between the inactive (yellow) and transitional (green) activity states assigned by the Hidden Markov Model.

**Additional file 3 - Linear mixed-effects models of denning phenology across demographic groups**

To assess whether denning phenology differed among demographic groups, we used linear mixed-effects models fitted. Separate models were built for den entry and den exit dates (expressed as Julian day). In each model, demographic group and year were included as fixed effects, while individual identity was included as random intercept to account for repeated measures of the same individuals across years. Estimated marginal means (LS-means) were computed for each demographic group. Pairwise contrasts between reproductive status categories were then performed using Tukey’s post hoc tests to control for Type I error inflation associated with multiple comparisons (Tables A3.1 and A3.2; Figure A3.1).

PF = Pregnant female; F COY = Female with cubs-of-the-year; F Y = Female with yearlings; AF = Solitary adult female; AM = Adult male; SF = Solitary subadult female; SM = Solitary subadult male.

FGB COY = Female who gave birth during the winter and with cubs-of-the-year; FGB CL = Female who gave birth during the winter but lost her cubs; FGB CU = Female who gave birth during the winter but whose cubs’fate is unknow; F Y = Female with yearlings; F TYO = Female with two years old offsprings; AF = Solitary adult female; AM = Adult male; SF = Solitary subadult female; SM = Solitary subadult male

**Table A3.1.** Pairwise comparisons of adjusted means (EMMeans) for date of den entry across Scandinavian brown bear demographic groups, with Tukey’s adjustment for multiple testing.

| Comparison | Estimate | SE | df | lower.CL | upper.CL | t.ratio | p.value |
| --- | --- | --- | --- | --- | --- | --- | --- |
| AM – F COY | 3.200 | 2.902 | 234.0 | -5.430 | 11.830 | 1.103 | 0.9268 |
| AM – F Y | 1.294 | 3.576 | 317.4 | -9.316 | 11.904 | 0.362 | 0.9998 |
| AM – PF | 13.143 | 2.558 | 176.1 | 5.514 | 20.772 | 5.138 | **< 0.0001** |
| AM – AF | 6.247 | 3.173 | 271.0 | -3.178 | 15.672 | 1.969 | 0.4370 |
| AM – SF | -4.043 | 2.783 | 220.7 | -12.325 | 4.240 | -1.452 | 0.7724 |
| AM – SM | -3.043 | 3.590 | 288.3 | -13.703 | 7.617 | -0.848 | 0.9796 |
| F COY – F Y | -1.906 | 2.976 | 252.0 | -10.751 | 6.938 | -0.641 | 0.9954 |
| F COY – PF | 9.943 | 1.776 | 258.2 | 4.665 | 15.220 | 5.599 | **< 0.0001** |
| F COY – AF | 3.047 | 2.644 | 317.1 | -4.800 | 10.893 | 1.152 | 0.9111 |
| F COY – SF | -7.243 | 2.223 | 340.3 | -13.836 | -0.650 | -3.258 | **0.0209** |
| F COY – SM | -6.243 | 3.454 | 283.6 | -16.500 | 4.014 | -1.807 | 0.5441 |
| F Y – PF | 11.849 | 2.748 | 262.3 | 3.683 | 20.015 | 4.311 | **0.0005** |
| F Y – AF | 4.953 | 3.394 | 301.7 | -5.123 | 15.028 | 1.459 | 0.7687 |
| F Y – SF | -5.336 | 3.089 | 315.9 | -14.502 | 3.829 | -1.728 | 0.5979 |
| F Y – SM | -4.337 | 4.035 | 329.0 | -16.307 | 7.633 | -1.075 | 0.9351 |
| PF – AF | -6.896 | 2.330 | 325.1 | -13.808 | 0.016 | -2.960 | 0.0510 |
| PF – SF | -17.185 | 1.817 | 341.0 | -22.575 | -11.796 | -9.458 | **< 0.0001** |
| PF – SM | -16.186 | 3.166 | 253.7 | -25.596 | -6.775 | -5.112 | **< 0.0001** |
| AF – SF | -10.289 | 2.562 | 325.0 | -17.891 | -2.687 | -4.016 | **0.0014** |
| AF – SM | -9.289 | 3.681 | 305.0 | -20.216 | 1.637 | -2.523 | 0.1546 |
| SF – SM | 1.000 | 3.353 | 279.0 | -8.959 | 10.959 | 0.298 | 0.9999 |

PF = Pregnant female; F COY = Female with cubs-of-the-year; F Y = Female with yearlings; AF = Solitary adult female; AM = Adult male; SF =

**Table A3.2.** Pairwise comparisons of adjusted means (EMMeans) for date of den exit across Scandinavian brown bear demographic groups, with Tukey’s adjustment for multiple testing.

| Comparison | Estimate | SE | df | lower.CL | upper.CL | t.ratio | p.value |
| --- | --- | --- | --- | --- | --- | --- | --- |
| AM – FGB CL | -2.752 | 3.663 | 282.1 | -5.430 | 11.830 | -0.751 | 0.9980 |
| AM – FGB CU | -14.134 | 3.190 | 238.2 | -9.316 | 11.904 | -4.431 | **0.0005** |
| AM – FGB COY | -21.535 | 2.591 | 183.6 | 5.514 | 20.772 | -8.312 | **< 0.0001** |
| AM – F TYO | -10.316 | 3.586 | 291.8 | -3.178 | 15.672 | -2.877 | 0.0987 |
| AM – F Y | -16.588 | 2.884 | 225.9 | -12.325 | 4.240 | -5.753 | **< 0.0001** |
| AM – AF | -9.680 | 3.152 | 253.6 | -13.703 | 7.617 | -3.071 | 0.0589 |
| AM – SF | -7.048 | 2.947 | 238.8 | -10.751 | 6.938 | -2.392 | 0.2933 |
| AM – SM | -8.825 | 3.772 | 286.9 | 4.665 | 15.220 | -2.340 | 0.3221 |
| FGB CL – FGB CU | -11.383 | 3.677 | 311.6 | -4.800 | 10.893 | -3.096 | 0.0542 |
| FGB CL – FGB COY | -18.783 | 3.096 | 309.0 | -13.836 | -0.650 | -6.066 | **< 0.0001** |
| FGB CL – F TYO | -7.564 | 3.903 | 291.4 | -16.500 | 4.014 | -1.938 | 0.5877 |
| FGB CL – F Y | -13.836 | 3.251 | 306.6 | 3.683 | 20.015 | -4.256 | **0.0009** |
| FGB CL – AF | -6.929 | 3.546 | 305.9 | -5.123 | 15.028 | -1.954 | 0.5767 |
| FGB CL – SF | -4.296 | 3.313 | 302.7 | -14.502 | 3.829 | -1.297 | 0.9317 |
| FGB CL – SM | -6.074 | 4.324 | 299.9 | -16.307 | 7.633 | -1.405 | 0.8955 |
| FGB CU – FGB COY | -7.400 | 2.564 | 309.6 | -13.808 | 0.016 | -2.886 | 0.0961 |
| FGB CU – F TYO | 3.818 | 3.582 | 307.9 | -22.575 | -11.796 | 1.066 | 0.9786 |
| FGB CU – F Y | -2.453 | 2.899 | 312.0 | -25.596 | -6.775 | -0.846 | 0.9953 |
| FGB CU – AF | 4.454 | 3.145 | 311.8 | -17.891 | -2.687 | 1.416 | 0.8909 |
| FGB CU – SF | 7.086 | 2.940 | 311.6 | -20.216 | 1.637 | 2.410 | 0.2822 |
| FGB CU – SM | 5.309 | 3.920 | 282.7 | -8.959 | 10.959 | 1.354 | 0.9136 |
| FGB COY – F TYO | 11.218 | 2.897 | 258.5 | lower.CL | upper.CL | 3.872 | **0.0043** |
| FGB COY – F Y | 4.947 | 1.939 | 244.3 | -5.430 | 11.830 | 2.552 | 0.2130 |
| FGB COY - AF | 11.854 | 2.489 | 311.0 | -9.316 | 11.904 | 4.764 | **0.0001** |
| FGB COY - SF | 14.486 | 2.273 | 309.3 | 5.514 | 20.772 | 6.373 | **< 0.0001** |
| FGB COY - SM | 12.709 | 3.456 | 266.2 | -3.178 | 15.672 | 3.678 | **0.0086** |
| F TYO – F Y | -6.272 | 3.101 | 253.6 | -12.325 | 4.240 | -2.022 | 0.5292 |
| F TYO – AF | 0.636 | 3.492 | 296.9 | -13.703 | 7.617 | 0.182 | 1.0000 |
| F TYO – SF | 3.268 | 3.350 | 304.4 | -10.751 | 6.938 | 0.976 | 0.9878 |
| F TYO – SM | 1.491 | 4.255 | 304.7 | 4.665 | 15.220 | 0.350 | 1.0000 |
| F Y – AF | 6.907 | 2.731 | 307.3 | -4.800 | 10.893 | 2.529 | 0.2223 |
| F Y – SF | 9.540 | 2.527 | 311.1 | -13.836 | -0.650 | 3.776 | **0.0059** |
| F Y – SM | 7.762 | 3.689 | 280.4 | -16.500 | 4.014 | 2.104 | 0.4724 |
| AF – SF | 2.632 | 2.844 | 309.3 | 3.683 | 20.015 | 0.925 | 0.9914 |
| AF – SM | 0.855 | 3.895 | 291.1 | -5.123 | 15.028 | 0.220 | 1.0000 |
| SF – SM | -1.777 | 3.735 | 286.8 | -14.502 | 3.829 | -0.476 | 0.9999 |

**Figure A3.1.**
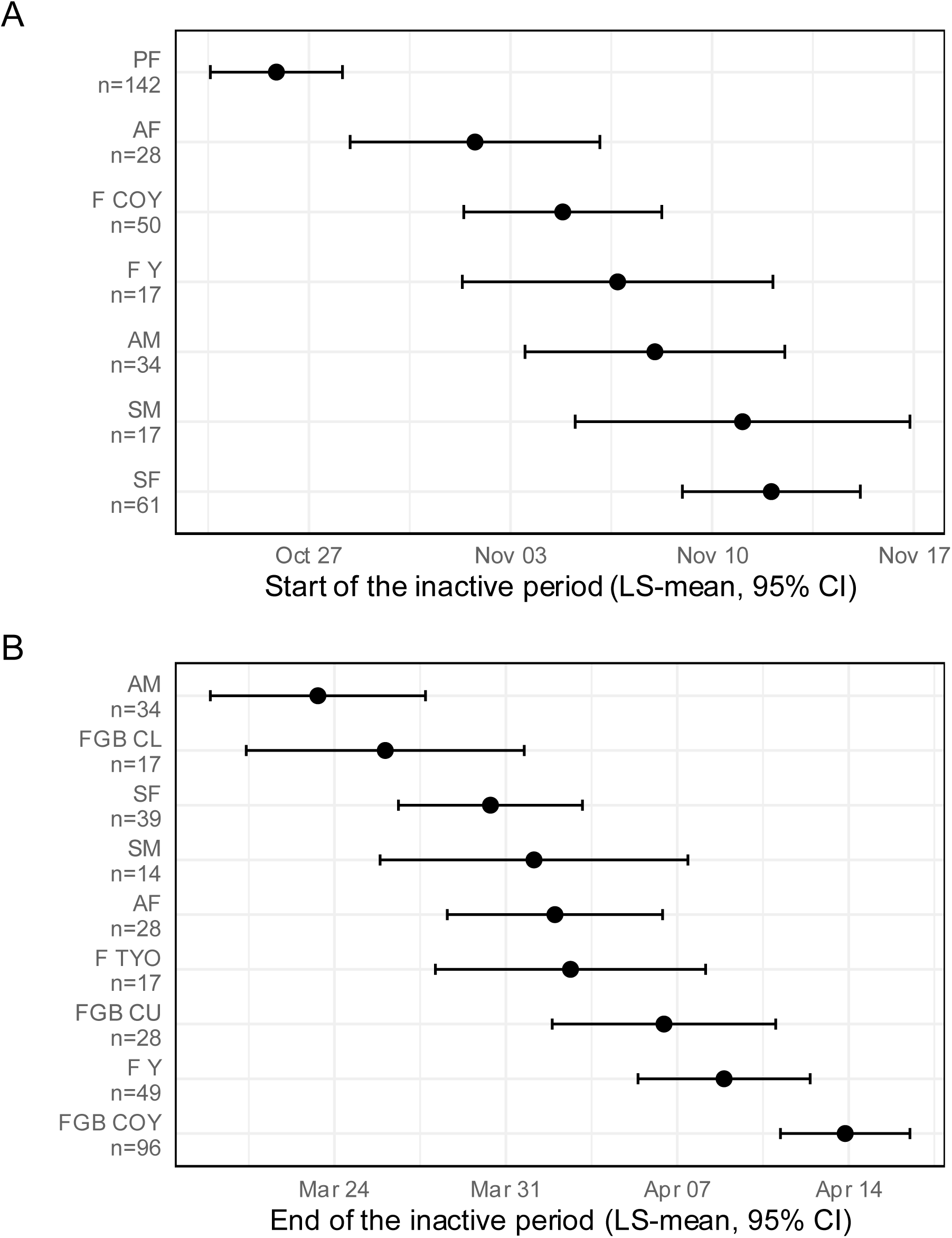
Plot of start (A) and end (B) LS-mean dates of the inactivity period by demographic group for Scandinavian brown bears in south-central Sweden between 2003 and 2024. PF = Pregnant female; F COY = Female with cubs-of-the-year; F Y = Female with yearlings; F TYO = Female with two years old offsprings; AF = Solitary adult female; AM = Adult male; SF = Solitary subadult female; SM = Solitary subadult male; FGB COY = Female who gave birth during the winter and with cubs of-the-year; FGB CL = Female who gave birth during the winter but lost her cubs; FGB CU = Female who gave birth during the winter but whose the fate of the cubs is unknow.

